# The Drosophila ovary produces two waves of adult follicles and a novel pupal wave that turns over

**DOI:** 10.1101/2025.11.11.687893

**Authors:** Wayne Yunpeng Fu, Allan C Spradling

## Abstract

Ovarian follicles in most species are assumed to develop using a single pathway. However, in Drosophila pupae, lineage tracing 2,937 single-cell clones identified three follicle “waves” that follow distinct programs arrayed anterior to posterior in the germarium and developing ovarioles. 40 primordial germ cells (PGCs) become anterior germline stem cells (GSCs) that produce “wave 2” follicles throughout adulthood. 100 PGCs posterior to wave 2 develop into “wave 1.5” follicles that form the earliest laid eggs. Different follicle stem cells (FSCs) sequentially occupy the same two niches timed to provide both waves with specific follicle cells, consistent with programmed differences between waves 2 and 1.5. “Wave 1” PGCs located even further posterior proliferate, form cysts, interact with swarm cells, degenerate, break from the ovary 22-26 hours after puparium formation, and release lipid-enriched vacuoles. Some testis germ cells behave similarly. We speculate that wave 1 germ cells contribute directly or indirectly to the pharate adult ecdysone pulse that mediates adult development and sex-specific neural remodeling. Why Drosophila follicle waves generally resemble those clarified recently in mouse preadult ovaries merits additional study.

## INTRODUCTION

Some species generate subclasses of ovarian follicles, in spatially or temporally separated groups known as waves (DeFalco and Capel, 2009; Eppig and Handel, 2012; Pazdernik and Schedl, 2013). Mammals such as mice generate wave 1 ovarian follicles that grow immediately at birth in the central medullar region, while the other ∼95%, wave 2, reside in the cortex region and become quiescent as the ovarian reserve (Mork et al., 2012; Rotgers et al., 2018). The fate of these waves was recently clarified and a third wave of primordial follicles, termed “wave 1.5,” was identified (Yin and Spradling, 2025). Wave 1.5 develops at the medulla/cortex boundary, arrests for ∼ 2 weeks, and produces most of the earliest ovulated oocytes rather than wave 1 (Meinsohn et al., 2021; Yin and Spradling, 2025). Wave 1 follicles generate few if any mature oocytes, but undergo a process classically termed “atresia” (Byskov, 1974). Studies of wave 1 atresia using lineage analysis, tissue clearing and 3D microscopy found that most oocytes and granulosa cells turn over. However, thecal cells proliferate, interstitial gland cells are recruited and these hormone-producing cells reorganize into a gland-like region that synthesizes mainly androgenic steroids during peri-puberty (Yin and Spradling, 2025), a sensitive period of embryonic development (Trova et al., 2021).

In Drosophila adults, ovarian follicles develop in ordered strings known as ovarioles that begin with a germarium region containing stem cells producing both germline and somatic cells (Figure 1B). 2-3 GSCs reside in a niche consisting of terminal filament (TF) and cap cells (CC) at the germarium tip where they give rise to daughter cells called cystoblasts (CB) that undergo 4 rounds of cyst division to form 16-cell germline cysts consisting of one oocyte and 15 nurse cells (Figure 1C, 1E). Two follicle stem cells (FSCs) reside midway through the germarium on opposite sides at the junction between regions 2a and 2b where cysts switch from a temporary covering of squamous escort cells (ECs) to a monolayer of epithelial follicle cells (Margolis and Spradling, 1995). Both FSCs are maintained in niches at the 2a/2b junction by local escort cells and intercellular signals (Nystul and Spradling, 2007; Sahai-Hernandez and Nystul, 2013).

**Figure 1.**
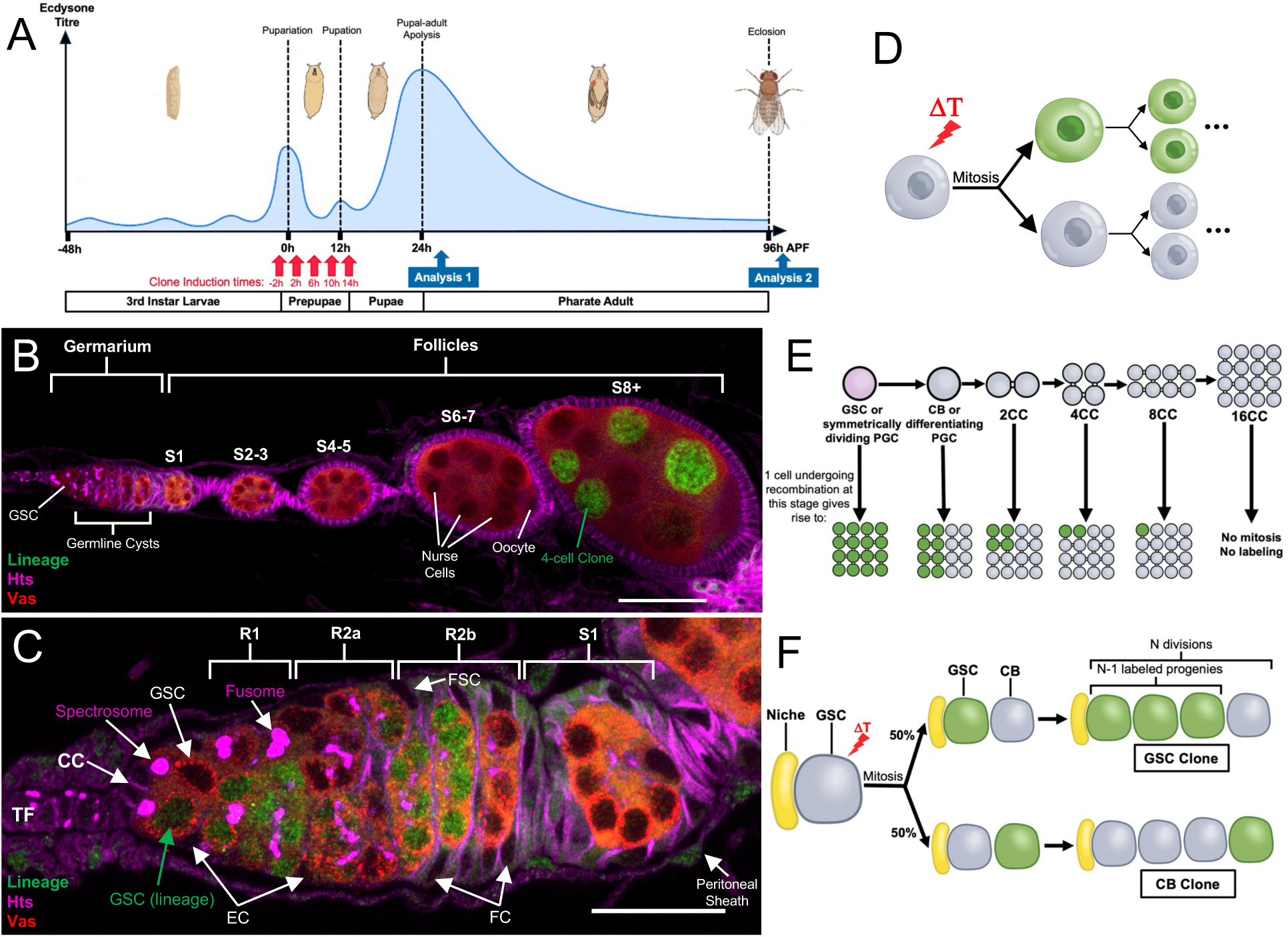
Ovary anatomy and strategy for analyzing ovarian follicle waves. **A.** Timeline of Drosophila development in hours after puparium formation (APF) and ecdysone levels during the third larval instar and pupal stages (modified from Scanlan et al., 2024). Red arrows show five clonal induction times; blue boxes show times of clone analysis. **B.** An ovariole from a fly heat-shocked at 2h APF is tipped with a germarium and 4 finished follicles. Its stage 8 follicle (S8) contains a clone labeling 4 of 16 germ cells. **C.** Germarium from a newly-eclosed fly heat-shocked at 2h APF. One lineage-labeled GSC and 4 progeny cysts are indicated. TF: terminal filament, CC: cap cells, EC: escort cells, FSC: follicle stem cell, FC: follicle cells. Fusomes: magenta (Hts). R1, R2a, R2b: germarium regions; S1: budding follicle. Scale bars: 50μm (B); 20μm (C.) **D.** Lineage labeling system. Heat shock (ΔT) activates a constitutive marker gene (green) in one daughter cell following mitosis (see Figure S1A.) **E.** Upper: Diagram of germline cyst formation from GSC, CB (cystoblast), 2-16-cell germline cysts (n-CC). Lower: clone size (green) following recombination at the indicated stages. **F.** Clone patterns induced by heat shock (ΔT) in a GSC. In GSC clone (above), N-1 daughter cysts are labeled; CB clone (below), the first daughter only is labeled.

The Drosophila ovary develops under control of the steroid hormone ecdysone during larval and pupal stages (Figure 1A; King et al., 1968, King, 1970; Sahut-Barnola et al., 1995; Godt and Laski, 1995; Forbes et al., 1996; Couderc et al., 2002; Camara et al., 2019; Reilein et al. 2021). The embryonic ovary acquires primordial germ cells (PGCs) which proliferate throughout larval development in association with somatic intermingled cells (Wieschaus and Szabad, 1979; Zhu and Xie, 2003; Li et al., 2003; Gilboa and Lehmann, 2006; Ganz et al., 2011; Banish et al., 2021; Rosales-Nieves et al., 2025). A pulse of the steroid hormone ecdysone induces the formation of TFs in third instar larval ovaries that guide ovariole development (Godt and Laski, 1995; Sahut-Barnola et al, 1996; Forbes et al., 1996). Before TFs are completed, a large population of somatic swarm cells, whose function is incompletely understood, migrate from the anterior to the posterior region of the ovary (Couderc et al., 2002; Banish et al., 2021).

Another key event is GSC niche formation (Lin and Spradling, 1993; Xie and Spradling, 2000; Zhu and Xie, 2003; Song et al., 2002; Song et al., 2007; Gilboa and Lehmann, 2006; Panchal et al., 2017). The TF constitutes the distal part of the GSC niche, whose Notch-dependent development continues during early pupal stages by adding CCs and specialized basement membranes (Zhu and Xie, 2003; Song et al., 2002; Song et al., 2007; Panchal et al., 2017; Yatsenko and Shcherbata, 2018; Diaz-Torres et al., 2021). 30-40 PGCs per ovary enter stem cell niches and become GSCs, but the timing of niche completion and GSC activation remain imprecise. About 60 other PGCs directly undergo oogenesis to generate the first ovulated eggs (Zhu and Xie, 2003; Asaoka and Lin, 2004), but their development has not been fully characterized.

Here we use lineage tracing and developmental studies to identify and characterize three Drosophila follicular waves, which we named based on their resemblance to the three waves of mouse follicles. “Wave 2” follicles are generated from GSCs and follicle cells generated by two adult-like FSCs. “Wave 1.5” follicles are generated from individual PGCs that form cysts and acquire follicle cells from two pupal-specific FSCs; they produce the first ∼100 eggs. A previously undescribed wave of follicles we term “wave 1” develops first but turns over between 22h and 26h after puparium formation (APF) via an atresia-like process. Wave 1 germ cells break open and release lipophilic cytoplasmic material, before they exit the ovary posterior along with somatic cells. Similar germ cell turnover in the pupal testis takes place at this time. We propose that wave 1 atretic gametes contribute in a currently undetermined manner to the pharate adult ecdysone pulse.

## RESULTS

### Analyzing ovarian follicle waves by lineage tracing

To characterize follicle waves, we used a background-free lineage labeling system based on heat shock-induced FLP-mediated recombination between chromosome 2L homologs (Harrison and Perrimon, 1994; Margolis and Spradling, 1995; Figure S1A). Heat shock can potentially activate recombination in any actively cycling cell but does so at low frequency, allowing for single-cell resolution. Recombination causes only one of the two daughter cells at the next mitosis to become heritably marked (Figure 1D), a property that aides analysis of germline development due to its invariant GSC and germline cyst division patterns (Figure 1E, F).

Clone induction was accurately timed (±2h) by using the short transition period into pupariation as a benchmark (see Methods). We induced clones at 4-hour intervals between -2h and 14h (all times refer to APF) and analyzed them either 24h later or shortly after adult eclosion (Figure 1A). All ovarioles were analyzed for somatic cell clones in both germaria and follicles, whereas we found it sufficient to analyze the larger number of germ cells in fewer ovaries. Altogether, we fully characterized 2,368 germ cell clones and 569 somatic cell clones from 2,270 individual ovarioles (Table S1-S3).

### Characterization of wave 2 follicles

We define “wave 2” follicles as those derived from GSCs, and “wave 1.5” as those produced from directly differentiating PGCs. First, we analyzed the GSCs that support wave 2 follicle production throughout adult stage. GSC clones can be recognized in newly eclosed adult ovaries since they contain a marked GSC at the tip of the germarium followed by adjacent marked progeny cysts resulting from GSC asymmetric divisions (Figure 1C, 2A). The number of GSC clones with these properties increased significantly at 10-14h (N=388) compared to the three earlier times ‘(N=1579, (p<0.001, Fisher’s exact test) to reach a labeling frequency similar to that of the adult GSCs (Figure 2C, blue line). Since lineage marker recombination only requires an active, post-G1 cell cycle state, the labeling rate of a cell population positively correlates with cell number, but not cell cycle length. Therefore, the increased labeling rate at 10h indicates a larger GSC precursor population. Marked GSCs produce an average of 4 marked progeny cysts by eclosion, and since their first division product is unmarked (Figure 1F), pupal GSCs usually undergo 5 divisions during pupal development (Figure 2C, orange line). Using ovarioles with all GSCs labeled we verified that wave 2 follicles remain in the germarium at eclosion (Figure 2D-D’).

**Figure 2.**
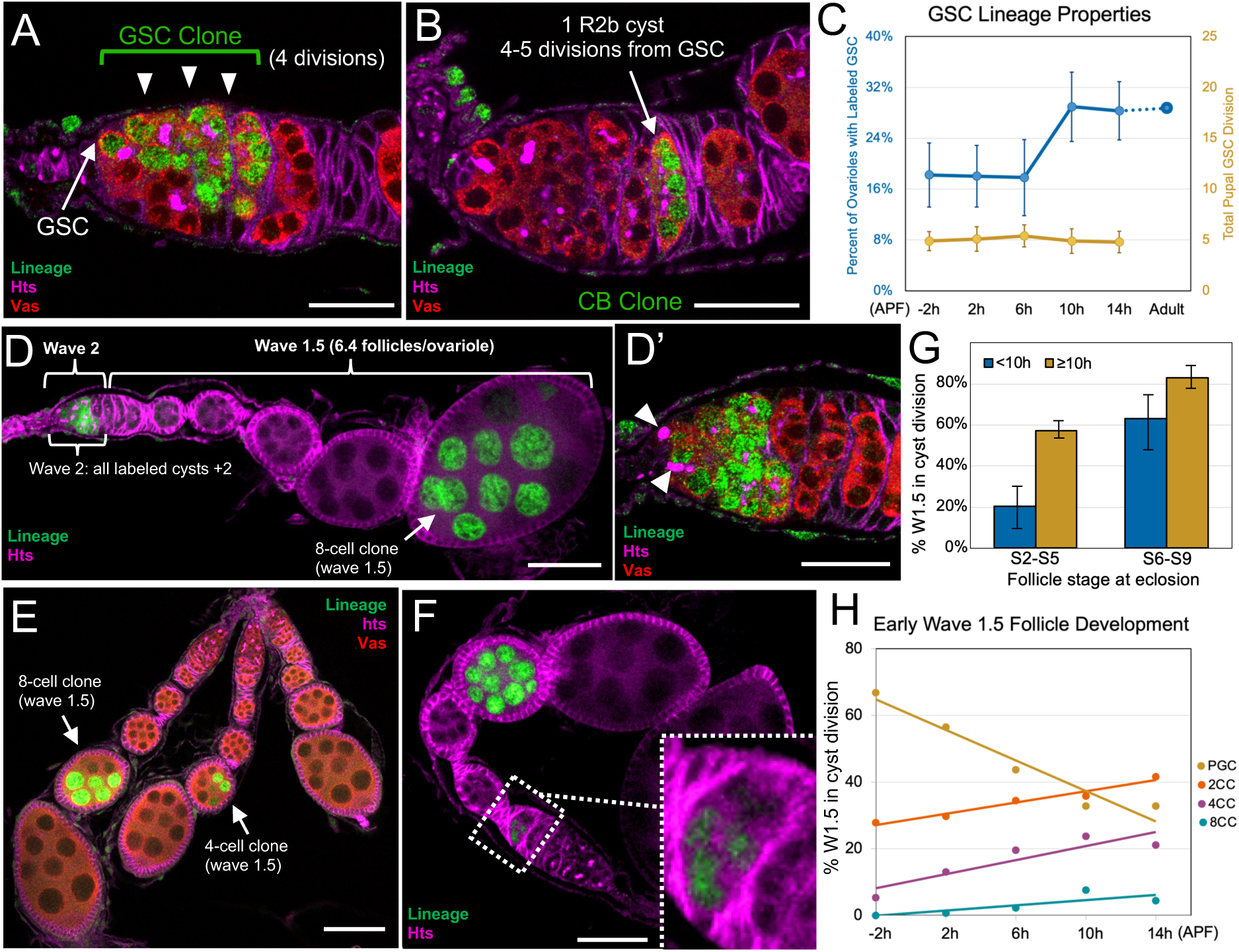
Identifying wave 2 and wave 1.5 follicles. **A.** GSC clone: LacZ-labeled GSC and 3 progeny cysts (arrowheads). **B.** CB clone: ovariole with single labeled 16-CC in R2B. **C.** GSC clone (blue) frequency (% ovarioles) vs induction time (hours APF). Number of GSC divisions (progeny cysts) in clone (orange). Ovariole numbers (N) =346, 303, 129, 224, 164 respectively. Mean +/- s.e.m. Adult value from Margolis and Spradling, 1995. **D-D’.** Ovariole with all GSCs labeled, showing location of wave 2 and wave 1.5. D’: Germarium region from D; all wave 2 (w2) cysts including 2 unlabeled cysts remain in germarium. **E.** Ovarioles with labeled w1.5 clones in germ cells. **F.** Ovariole with fully labeled w1.5 follicle at S4-5 due to PGC symmetric division; a w1.5 follicle at S1 is magnified at right. **G.** Percentage of wave 1.5 follicles that were no longer PGCs at induction time points before 10h (blue) vs after 10h (tan), in young vs older follicles at eclosion (Stage 2-5 vs Stage 6-9). mean +/- s.e.m. across time points in each group. **H.** Percentages of wave 1.5 follicles PGCs and cyst stages versus times of clone induction showing progression of cyst divisions. Sample size for 2G and 2H are the same as in 2C. Scale bars are 20μm in A, B, D’, 50μm in D, F, G.

A second class of GSC-related clones was termed CB clones (Figure 2B). CB clones were recognized as single 16-cell labeled cysts located ∼5 divisions downstream from unlabeled GSCs. They are predicted to result from segregation of the lineage marker gene into the CB daughter rather than the GSC daughter at the first division of a recombined GSC (Figure 1F). Across all time points, we observed 199 CB clones and 221 GSC clones. The closeness of these numbers to 50:50 segregation limits the role of GSC symmetric division plays in filling early niches. As predicted, CB clones were located on average more distal than the oldest cyst in marked GSC clones.

### Characterization of direct-developing wave 1.5 follicles providing early fertility

The most common clonal pattern was partial labeling of 8 or fewer germ cells in a single follicle. Such clones result from recombination within a direct developing PGC or a downstream cyst cell, and therefore mark wave 1.5 follicles (Figure 1E, 2E; Table S3). Rarely, wave 1.5 follicles contained 16-cell clones due to prior symmetric division of a recombined PGC (Figure S2B) before it began direct cyst formation (Figure 2F). Marked wave 1.5 cysts were also observed in germarium region 2b (Figure 2F) and S1, posterior to the oldest wave 2 cysts, as expected.

Comparing the labeling pattern of wave 1.5 follicles across time points, we found that the youngest wave 1.5 follicles preferentially initiate cyst division coincident with the 10h ecdysone pulse, whereas the two most mature follicles have already undergone more cyst divisions and responded less (Figure 2G). Averaging all wave 1.5 follicles at each time point, a steady decline in the percentage of undivided PGCs (8-cell clones) is seen as a function of clone induction time (Figure 2H). Correspondingly, there is a steady increase in the percentage of cysts undergoing successively later rounds of cyst division. These data show that wave 1.5 cyst divisions proceed at a similar average speed, with 67% remaining as undivided PGCs at -2h whereas only 33% of clones mark PGCs that have not yet begun cyst divisions at 14h.

Since all germline cysts and follicles within ovarioles at eclosion are derived from either GSCs (wave 2) or direct-developing PGCs (wave 1.5), we quantified wave size. On average, each ovariole contains 12.5 wave 2 cysts (5 ±0.9 cysts produced per GSC and 2.5 ±0.6 GSCs per ovariole, n=165 and ranges represent SD). By subtracting wave 2 cysts from the total 19 ±2.4 cysts and follicles present in an ovariole at eclosion, it was determined that wave 1.5 was responsible for about 6.5 follicles per ovariole (Figure 2D). Thus, since an ovary contains 15.5 ±1.1 ovarioles, ∼100 wave 1.5 follicles are produced per ovary, a much larger population than previously estimated.

### A large, novel population of wave 1 germ cells turns over between 22h-26h APF

Direct examination of pupal ovaries from -2h to 30h APF allowed us to further study developing wave 2 and wave 1.5 follicles, and revealed an unexpected third follicle wave that we term “wave 1” (Figure 3A-J). The number of single germ cells plus cysts in each ovariole was determined based on fusome morphology beginning at 6h (Movie S2). It continues to increase as some GSCs undergo symmetric division and ovarioles incorporate all germ cells within the sheath (Figure 3D) until 18h (Figure 3K).

**Figure 3.**
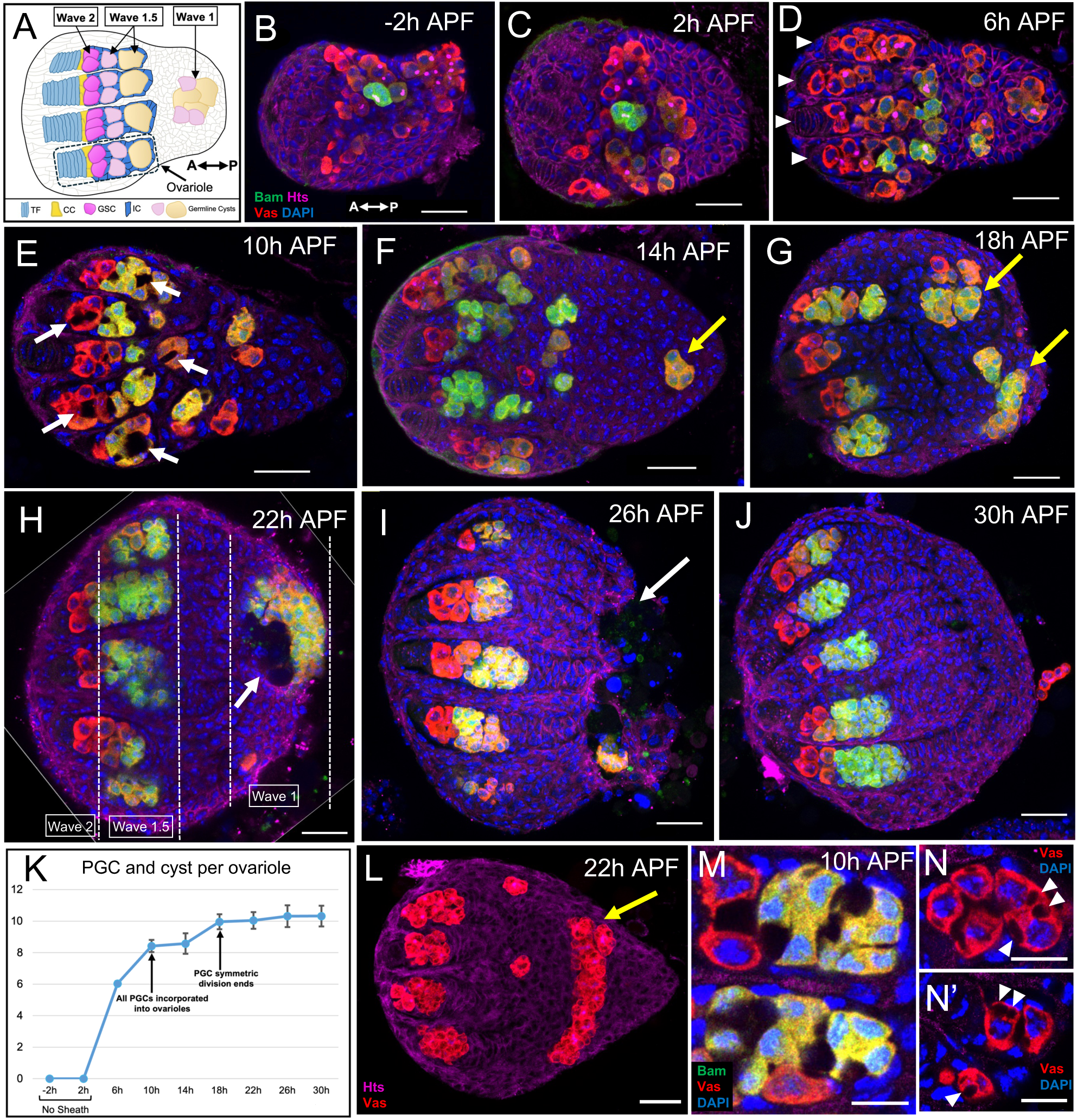
Time course of pupal germ cell development in three follicle waves. **A.** Schematic of a 10h APF pupal ovary oriented as in the following images, anterior to the left and posterior to the right. Arrows on top point to PGCs that become each of the three waves. **B-J.** A time sequence of pupal ovaries from -2h APF to 30h APF stained for germ cells (Vasa, red), forming cysts (bam-GFP, green), fusomes and somatic cell membranes (Hts/1B1, magenta) and nuclei (DAPI, blue). Arrowheads in D indicate newly formed ovarioles where ovariolar germ cells are separated by the sheath. Yellow arrows in F-G indicate the initial posterior wave 1 germ cell precursors. White arrows in E, H, I point to vacuoles devoid of antibody labeling that appear around wave 1 germ cells. The distinction of 3 waves is outlined in H. **K.** Quantification of total single PGCs and cysts per ovariole from -2h to 30h APF. Error bars represent SD across ovaries. n=10 ovaries for each time point. **L.** Stacked image of a 10μm section to show the large aggregation of all wave 1 germ cells into a disk-shaped mass in the posterior (arrow) of a 22h APF ovary. **M-N.** 10h APF germ cells magnified to show vacuoles and gaps between cells. Arrowheads point to the vacuoles in N-N’ inside germ cell cytoplasm. Scale bars are 50μm in B-J and L, 10μm in M-N, N’.

Beginning as early as 6h (Figure 3D), a subpopulation of PGCs migrates toward the posterior of the ovary. Posterior somatic cells form a conical structure at this time (Figure 3E-F, L). Germ cell posterior movement continues, producing a distinct group (Figure 3F-G, L, arrows). The germ cells at the posterior further increase in number by forming cysts and by 22h they aggregate into a mass of approximately 150 cells (Figure 3G-H, L; Figure 4D, orange line). Between 22-26h APF, all the posterior germ cells and some adjacent somatic cells are extruded from the ovary posterior (Figure 3I). We term these cyst structures “wave 1” because of their early development and atresia-like turnover, akin to wave 1 follicles in the mouse. All three waves are visible along the A-P axis in Figure 3H. By 30h, the ovary has completely lost wave 1 germ cells and its protruding posterior has been repaired, appearing spherical in shape. (Figure 3J).

**Figure 4.**
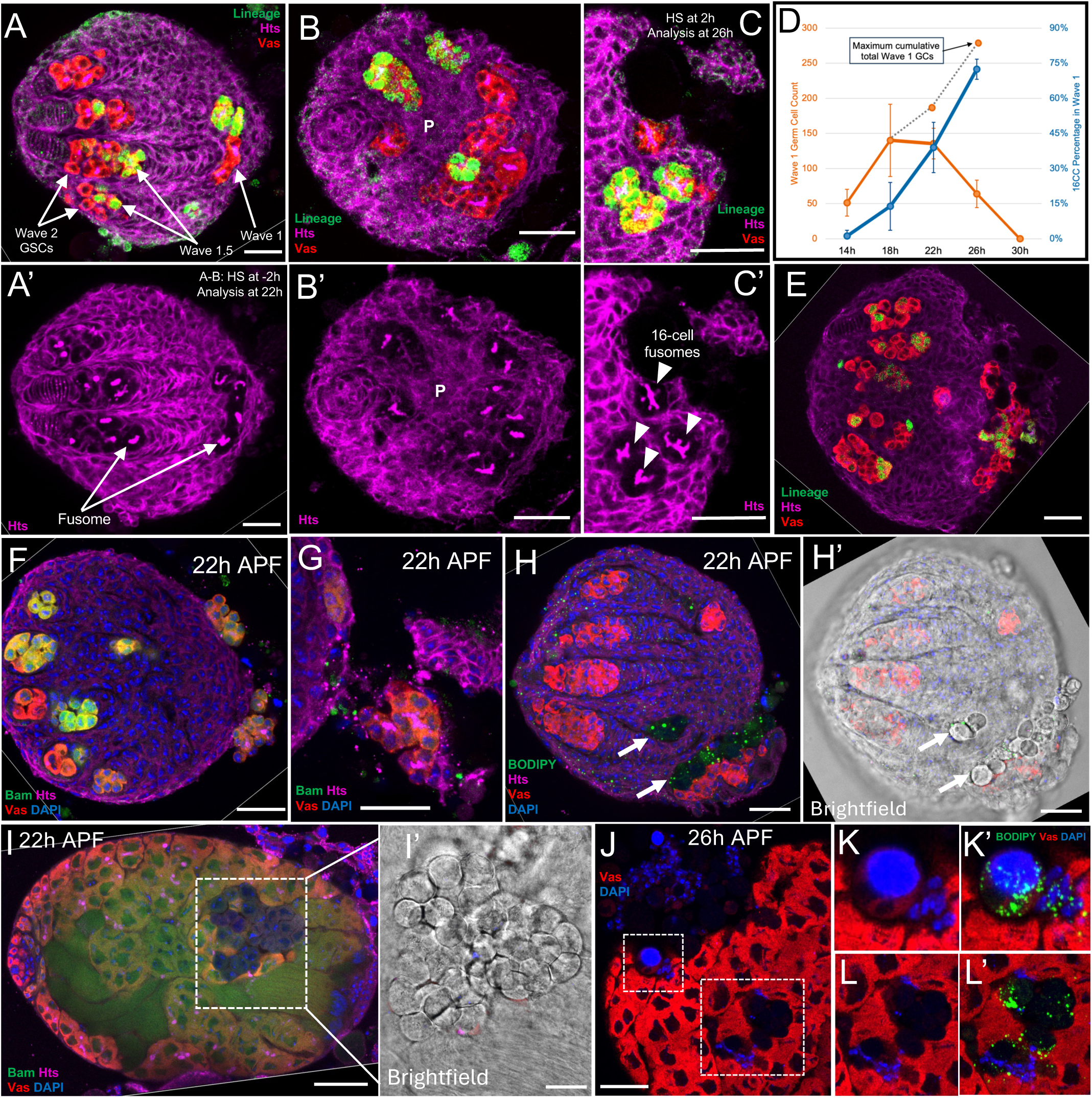
Rapid proliferation of ovarian wave 1 germ cells and cysts. **A-A’.** Pupal ovary lineage labeled at -2h APF and analyzed at 22h APF. Wave 1 germ cells at the posterior include lineage-labeled cysts with fusomes (A’, arrows). **B-B’.** An ovary with the same conditions as in 2A viewed from the posterior to reveal 14 wave 1 cysts, all with fusome (B’); 4 cysts are fully labeled. **C-C’.** Pupal ovary labeled at 2h APF and analyzed at 26h APF. A large segment of ovarian posterior region has broken off (upper right). C’ shows that the cysts all have 16-cell fusomes, 3 are lineage marked in all cells. **D.** Quantification of wave 1 germ cells (orange) from the time they start to become spatially separated from the other two anterior waves (14h APF) until the time they have all been extruded from the ovary (30h APF). Points on dashed lines are cumulative total wave 1 germ cells ever produced discounting loss from extrusion and considering the continuous cyst amplification, as shown by the constant increase of 16 cell-cyst percentage out of all cysts in wave 1 (blue). n=10,12,12,9 ovaries, respectively. Error bars represent SD across ovaries. **E.** Pupal ovary labeled at 2h APF and analyzed at 26h APF showing significant germ cell extrusion from the posterior, among which two large clusters are lineage positive. **F-G.** 22h APF bamP-GFP fly line pupal ovaries showing germ cell extrusion similar to that seen in lineage tracing fly line ovaries. **H-H’.** 22h APF ovary with lipid-enriched vacuoles stained with BODIPY embedded in the ovary next to wave 1 germ cells. H’ is the brightfield image of the same ovary showing morphology of the vacuoles (arrows). **I-I’.** A 22h APF testis with a large cluster of dying germ cells. I’ shows the morphological change of this germ cell cluster under brightfield. **J.** A 22h APF testis showing significant amount of cell loss on the upper left corner. Areas in squares are magnified in K and L. **K-L.** Dying testical germ cells with remnant vasa expression. K’ and L’ show lipid enrichment similar to that of 4H. Scale bars are 50μm in all images.

In 10h ovaries, clear vacuoles appear around germ cells (Figure 3E, M, N), coincident with the small prepupal ecdysone pulse (Figure 1A). Smaller vacuoles were observed inside germ cells, confirming germ cells to be the origin for such vacuoles (Figure 3N-N’). These vacuoles disappeared within 4 hours without causing lasting morphological changes to the tissue (Figure 3F).

### Rapid proliferation and clearance of wave 1 germ cells

We analyzed lineage-labeled germ cells directly in pupal ovaries 24 hours after clone induction to gain further insight into wave 1 development. Analyzing clones induced at -2h and 2h in pupal ovaries 24 hours later showed that individual wave 1 clones typically contain multiple nearby cysts that were fully labeled (Figure 4A-C), indicating wave 1 precursor PGCs expand rapidly by symmetric division and then by forming germline cysts with typical fusomes. Hts staining reveals branched fusome structures that are diagnostic of cyst division stage. Viewing the ovary from the posterior reveals branched 8 or 16-cell fusomes in >10 remaining wave 1 cysts (Figure 4B-B’). Adjacent fully labeled cysts indicate their common origin from a single PGC labeled 24 hours previously (Figure 4C-C’). We estimated the cumulative total number of wave 1 germ cells released by first determining that at 18h APF when single wave 1 forming cells and cysts reach a maximum, there are 22 wave 1 forming units in various stages of cyst formation as indicated by fusome structure (Figure 4D, S2D, S2E). At this time before cells began to be extruded, only 15% of cysts contain 16 cells (Figure 4D, blue line). However, at 22h and 26h APF the percentage of 16-cell cysts among the remaining wave 1 cysts reached 75%. Thus, cyst divisions continue even as clusters of wave 1 germ cells and associated somatic cells break off in groups at the posterior (Figure 4D, E-G) lowering the remaining total. Taking into account apparent individual variation, and cell loss over a period of time, we estimate that 250-300 wave 1 germ cells are expelled, with a maximum of around 22 founder cells x 16 = 350.

Immediately before and as wave 1 germ cells exit the ovary, large spherical vacuoles with significant BODIPY-stainable lipid droplet enrichment appear around them, likely derived from these germ cells (Figure 4H-H’). We examined 22-26h APF testes as well, and found a similar sequence of regional germ cell appearance changes (Figure 4I-I’) followed by expulsion from the gonad (Figure 4J-L) with concurrent lipid droplet accumulation (Figure 4K’, L’). In the testis, however, cells were lost from the sides of the gonad rather than the posterior, where germ cells are absent. This indicates that germ cell extrusion is not a female-specific process, and may serve a more general purpose in development. The fact that wave 1 extrusion temporally overlaps the onset of the pharate adult ecdysone pulse, and includes a release of lipid-rich vacuoles, suggests a causal relationship between these events.

### Ovarioles contain two pupal FSCs that resemble adult FSCs

In normal adult ovarioles, two long-lived FSCs each stably associate (t_1/2_ ∼10 days) with a separate niche located on opposite sides of the germarium at the 2a/2b junction. Adult FSCs divide each supply one founder follicle cell to each passing germline cyst shortly after they lose their covering of escort cell membranes. Each founder cell continues dividing to produce 50% of the continuous follicular monolayer comprising 60-80 cells in about 48 hours, that is sufficient for follicle budding, including polar and stalk follicle cells. Electron microscopy (King et al., 1968) and our observations (Figure S1C) show that cysts in early pupal ovarioles do not initially move side by side in a region 2a as in adults. This obscures the location of a possible 2a/2b junction, and FSC niche. Consequently, it was unclear where pupal FSCs are located or if follicle cells generated during pupal stage arise in a different manner than adults.

Lineage analysis of the the somatic cell clones we observed at eclosion, however, resolved this problem and revealed adult-like FSC clones (Figure 5A) that define a pupal FSC population (”FSC-P”). Initially, at stages -2h to 6h, these clones were relatively rare (3.6%, N=1,587), but at 10h APF they were observed significantly more frequently (Figure 5B; 14.3%, p<0.0001, Fishers exact test N=683). The longest FSC-P clones initiated at precisely the same position at the 2a/2b junction described for adult FSCs. Like adult FSC clones, follicle cells in FSC-P clones were labeled in largely contiguous patches on adjacent follicles. However, most FSC-P clones spanned a smaller number of adjacent follicles, and frequently did not extend all the way to the 2a/2b junction (Figure 5C-C’, D). This is the observed pattern of an adult FSC clone that initiated at the 2a/2b junction, but after some number of cysts had passed whose labeled FSC had been replaced by an unlabeled FSC. The unlabeled FSC did not disturb ongoing follicle production and posterior movement, with the result that the labeled group of follicles simply continued to develop and move down the ovariole.

**Figure 5.**
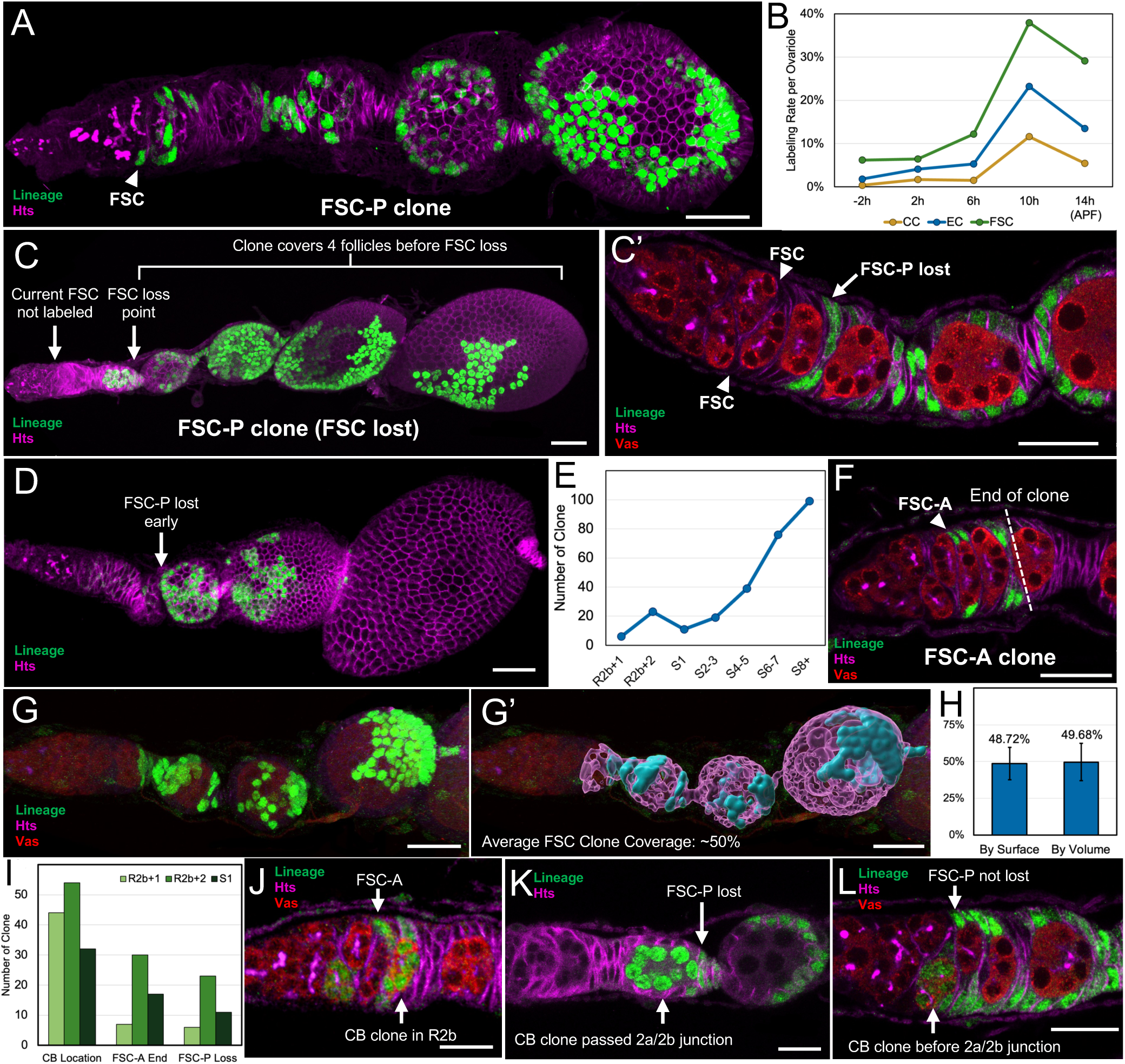
FSC development correlates to follicle waves. **A.** An FSC-P clone marked at 10h APF and analyzed by eclosion. Arrowhead points to the marked active FSC at 2a/2b junction. **B.** Somatic cell clone labeling rate as a function of induction times in hours APF. All cell types exhibit a significant increase in labeling rate at 10h (see text). While FSC clones maintained the elevated rate, that of cap cell (CC) and escort cell (EC) clones dropped subsequently at 14h. N=621, 517, 449, 224, 459 total scored ovarioles. **C.** An FSC-P clone without a labeled active FSC, where the labeled FSC-P was replaced by an unlabeled FSC after its progeny has covered 4 sequential wave 1.5 follicles. **C’.** Germarium containing an FSC-P clone that does not contain an active FSC at 2a/2b junction (arrowheads). **D.** An FSC-P clone that lost its FSC early, only covering two early wave 1.5 follicles. **E.** Distribution of FSC-P clone loss points by follicle position. n=273 clones scored. **F.** A short FSC-A clone labeled at 14h APF that by eclosion has only covered two R2b cysts. **G-G’.** Example demonstrating how FSC-P follicle coverage rate was calculated in Imaris. G shows the raw signal, G’ shows the clone coverage in blue and total FC layer on follicles with the clone in magenta. **H.** FSC-P follicle surface coverage rate calculated directly by surface area of the 3D objects created in Imaris, or by surface area converted from volume of such objects. n=26 follicles measured. **I.** Cyst location distribution of CB clones, FSC-A end points and FSC-P loss points. R2b+1 stands for the first R2b cyst position, etc. Only clones of these types that involve a position in R2b and S1 are recorded in the graph. n=130,54,40 clones for each clone type. **J.** Ovariole with an FSC-A clone surrounding a CB clone. **K.** Ovariole with a CB clone past the 2a/2b junction also containing an FSC-P clone that does not contain an active FSC, where the FSC-P loss point is immediately before the CB clone. **L.** Ovariole where a CB clone has not yet passed the 2a/2b junction also contains an FSC-P clone that has not lost its active FSC. Scale bars are 20μm in all images.

Interestingly, the frequency of FSC-P displacement was much higher overall than in adult FSCs which are half replaced about every 10 days (20 divisions) (Margolis and Spradling, 1995). We measured the number of replacement events at different follicle stages along the ovariole (which each correspond in temporal order to single FSC divisions). We recorded the start and end point of all FSC-P clones that spanned two or more follicles. Significantly more FSC-P loss events occurred on the most mature wave 1.5 follicles, usually reaching stage 8-9 at eclosion. Fewer losses occurred at each successively younger follicle, showing that initial FSC-P stability rapidly increases (Figure 5E). Each stability increase between stage 8+ and stage 2-3 follicles was significant (p<0.025, Fisher’s exact test, two tailed, N=332). Measurements of the follicle surface using 3D reconstruction showed that FSC-P clones from “lost” FSC-Ps still label an average of about 49% of follicle surface area (Figure 5G, G’, H; see also Movie S3-S4). Thus, the clones show that young FSC-P pairs do not deviate on average from depositing one cell each onto every cyst (except in the oldest wave 1.5 follicle, see below).

### The first wave 2 follicle activates a new set of FSCs

A second class of FSC clones was observed to arise much later in pupal development. They also initiate at the 2a/2b junction, but have only produced follicle cells for 1-3 subsequent cysts (Figure 5F). These were termed “FSC-A” (adult FSC) clones since the cysts they cover closely matched the location of the oldest wave 2 cysts, and they remained active following eclosion. To compare the relative location of FSC-A covered and wave 2 cysts, we measured the location of CB clones which mark initial wave 2 follicles and compared it with the most distal cysts marked by FSC-A clones. >75% of CB clones are found at the first or second region 2b cyst, and less frequently at the S1 follicle (Figure 5I). This indicates that the first wave 2 cyst passes the 2a/2b junction, and moves 1-3 additional cyst positions downstream. Notably, the average distribution of FSC-A clone end points closely matched the CB clone distribution (Figure 5I). In addition, the loss points of FSC-P clones that had most recently lost their FSC showed a very similar distribution to FSC-A and to CB clones (Figure 5C’, I). This suggests that loss of FSC-Ps in the germarium are often caused by FSC-A displacement, which may account for the small peak at “R2b+2” (second cyst position in R2b) in the FSC-P loss curve (Figure 5E).

To precisely correlate FSC-P replacement by FSC-A, we analyzed rare ovarioles in which both the first wave 2 cyst and one of the FSC types were lineage labeled. CB clones located at the first R2b position were co-labeled with FSC-A clones that did not extend beyond that cyst (Figure 5J). FSC-P clones were also observed to end immediately before the advent of a CB clone when these two types of clones are labeled in the same ovariole (Figure 5K); and when a CB clone has not yet reached the 2a/2b junction, the co-labeled FSC-P clone retained its active FSC at the 2a/2b junction (Figure 5L). These double clones strongly support the model that FSC-As replace FSC-Ps upon the advent of the first wave 2 cyst. As a consequence, FSC-A evidently produce the cells covering wave 2 follicles while FSC-Ps produce follicle cells on wave 1.5 follicles.

### Wave 1.5 follicles showed distinct properties

Wave 1.5 follicles showed an elevated frequency of cyst and follicle defects compared to normal adult follicles. Initial wave 1.5 follicles also had a higher frequency of germline cyst developmental abnormalities resulting in 8 or 32 cells per follicle (Figure S2H), indicating they underwent one less or one more round of cyst division. Such abnormalities were recorded in 3.8% (N=449) of the most mature wave 1.5 follicle, significantly more than the 1.6% (N=982) of stage 1-8 follicles (p< 0.025, Fisher’s exact test; see Table S3). In addition, we noticed an apparently elevated frequency of morphological defects, including oversized polar cells, suggesting increased polyploidization (Figure S2F). The number and size of border cell clusters, which include the anterior polar cells, were sometimes enlarged and delayed in migration (Figure S2F-G). Follicle cells on the first (oldest) wave 1.5 follicle in each ovariole displayed a distinctly shaped posterior patch on 68% of all clones at this location (Figure S3G-G’, N=72). Rather than coming from FSCs, these patches may have derived from basal stalk cells, which touch the posterior of the oldest wave 1.5 follicle and were previously reported to contribute to pupal follicle cells (Reilein et al. 2021). Similar follicle cell movement from interfollicular stalks onto the main body is known to take place in the adult ovarioles (Nystul and Spradling, 2010).

### Cap cell and escort cell differentiation correlates with niche development

Clones were also recorded of cap cells (CC) and specific escort cells (EC) clones (Figure 5B) that could be recognized by location (Figure S4A, K). EC1-3 locate around the GSC niche, and EC4-6 are close to FSC niches. An EC1 wraps the surface of each GSC, while EC2 contacts CBs (Figure S4B-C). A median EC3 is also present between the GSCs (Figure S4D). Three ECs similarly surround both FSC niches (Gao et al., 2019; Rust et al., 2020). EC4 is on the surface of the germarium and directly contacts the FSC (Figure S4E), more anterior to it’s EC5 partner (Figure S4F). Another median EC type, EC6, is found between the two FSCs located internal in the germarium (Figure S4G). At least in some cases, EC6 cell membranes extend to an adjacent FSC (Sahai-Hernandez and Nystul, 2013).

The labeling frequencies of both EC and CC clones increased significantly at 10h APF (Figure 5B, p<0.0001, Fishers exact test; N=1587). Since these clones were usually single cells, or in very small groups (Figure S4I), it suggests that ECs and CCs frequently undergo a terminal division(s) around 10h APF. Observed EC clone sizes confirm they sometimes share a common precursor, since 48% contain more than one EC, but 99% contain 4 or fewer cells. 84% of CC clones were single-cell clones, with the rest containing 2 cells (Table S2). The increased labeling rate at 10h was larger for EC1 and EC2, the most anterior ECs located next to the GSC niche, than for other ECs (Figure S4H). Clones of CCs, the other GSC niche cell, also showed a similar increase (Figure 5B), arguing that the GSC niches may finish production at this time.

## DISCUSSION

### Drosophila females produce three waves of follicles in a precise process

Our experiments demonstrated that the well-studied GSC-based system of follicle production in Drosophila (wave 2) is preceded by two earlier follicle waves. By accurately timing the many cellular and developmental steps in wave formation relative to 0h APF using lineage analysis, we gained a new understanding of when and how these waves arise. Wave 1.5 and wave 1 follicles directly derive from PGCs which expand by symmetric division to a final population of 160 PGCs by 18h APF (Figure S2A), a larger pool than previously realized. Many cap cells differentiate only at 10-14h APF (Figure 5B), suggesting that GSCs wait for niches to be completed before beginning to divide and produce new cysts after 10h. In contrast, 28% of wave 1.5 PGCs have already undergone 1, and 5.4% have already undergone 2 cyst divisions by 0h APF (Figure 2H). Indeed, most of pupal ovarian follicle development is centered on producing 100 wave 1.5 follicles. Most surprisingly, at 22-26h the posterior ∼50% of the pupal ovarian space, ∼20% of posterior somatic cells and ∼50% of pupal germ cells are occupied with generating wave 1 follicles, which were not previously known to exist due to their quick subsequent turnover.

### The ovary prioritizes rapid production of around 100 wave 1.5 follicles

Our studies argue that wave 1.5 follicles are intrinsically important, rather than just redundant products of excess PGCs needed to ensure that GSC niches are filled, as sometimes suggested earlier. All follicles that have formed by adult eclosion, as well as 1-2 posterior cysts in each germarium are of wave 1.5 origin. The need to produce these follicles in only about 5 days, compared to the 7-day developmental period of GSC-derived follicles (King, 1970; Lin and Spradling, 1993), likely explains precocious wave 1.5 cyst formation in late larval ovaries. Rapid production of wave 1.5 follicles might explain why they contained an elevated frequency of minor developmental defects. The defective follicles may not become fertile oocytes, however. In adults, cysts with other than 16 cells are rare and have never been shown to produce eggs; such cysts arising from mutation are generally sterile. Other more viable defects in wave 1.5 cysts may also exist. Despite this, wave 1.5 is the major activity of pupal ovarian development.

A strong suggestion that wave 1.5 follicles differ from wave 2 follicles comes from our finding that their follicular layer is usually produced by distinct stem cells, FSC-P, in contrast to wave 2 follicles, which are coated by FSC-A progeny. At present we lack transcriptomes to compare these two FSC types, nor have functional tests on laid eggs been performed. However, our finding here that pupal follicle layers are produced by two separate and coordinated stem cells, as in adults, implies that two replacement events are required to substitute FSC-A for FSC-P. FSC-P daughters cannot be responsible for the switch, because this would elevate the frequency of FSC-P clones that continue to cover wave 2 follicles even after wave 2 cysts begin to pass the region 2a/2b junction, or generate 100% follicle labeling, neither of which was observed. Indeed, it is difficult imagine how a programmed replacement would evolve if FSC-P and FSC-A were identical.

Wave 1.5-derived eggs probably provide some advantage to the great majority of Drosophila females, who in the wild live in resource-poor circumstances and never experience the abundant nutrition and oviposition sites needed to deplete these oocytes and begin laying wave 2 eggs. Wave 1.5 follicles are less dependent on adult nutrition than wave 2 follicles because they develop largely using larval nutrients stored in the fat body. Therefore, if FSC-P derived follicle cells synthesize more desiccation-resistant eggshells, it would allow more ovulation in drier, resource-poor microenvironments. In contrast, wave 2 follicles might produce eggs specialized for development and growth in moist, microbe-rich conditions on nutritious food sources in competition with other larvae.

### Drosophila wave 1 follicles may contribute to the pharate adult ecdysone pulse

Our finding that an early Drosophila germ cell wave produces 250-350 germ cells in germline cysts that turn over between 22h and 26h APF strongly suggests these wave 1 follicles have a function other than gametogenesis. Wave 1 germline cysts are extruded from the ovary together with a group of somatic cells that form a conical structure at the posterior (Figure 3L). Some of these may be swarm cells, a population that migrates from the anterior to the posterior during L3 (Couderec et al. 2002).

Our study shows that wave 1 cells are unlikely to represent larval tissue remodeling because they constitute a new cell population recently produced in a highly programmed manner that rapidly proliferates before collective turnover. Wave 1 extrusion occurs as the pharate adult ecdysone pulse (PAP) is rising, and ceases near its peak. While essential for pupal metamorphosis, the source of the PAP is currently unclear (Scanlan et al. 2023). Because the prothoracic gland turns over early in pupal development (Dai and Gilbert, 1991), these authors suggested that an unidentified tissue releasing ecdysone recycled and stored in an inactive from larval and/or prepupal development was the most likely source. Wave 1 germ cells in the ovary, and a similar subgroup of germ cells in the testis both turn over and release lipid-enriched vacuoles suggesting they promote a common purpose. Wave 1 follicles may themselves store and release ecdysone, signal other tissues to do so (Zhang et al. 2024), or both. Determining the true function of transient wave 1 follicles will ultimately require a separate study that is now possible based on their properties reported here.

### Drosophila pupal ovarian FSCs offer insight into stem cell niche development

The behavior of stem cell niches represent an important aspect of stem cell biology (Morrison and Spradling, 2008). We identified pupal FSCs and showed that precisely 2 FSC-Ps reside in niches comprised of cells located at the 2a/2b junction in newly eclosed ovarioles, the same number and location as adult FSCs. FSC-A stem cells arising in late pupae reside at this same location. Nothing resembling a 2a/2b junction has been identified in the larval or prepupal ovary but the differentiation of these cells at about 10h APF that likely involving EC4-6 cells (Figure S4H-I) would be a good place to begin investigation. The dramatic 5-fold increase in FSC-P stability in its niche during early wave 1.5 development (Figure 5E) also presents an opportunity to better understand factors involved in stem cell niche development. Pupal FSC niches are able to coordinate the division of two stem cells with passing cysts, resulting in the association of one daughter from each FSC per cyst even before they finalize full niche stability.

The programmed replacement of the two FSC-P stem cells in each ovariole by two new FSC-A stem cells associated with the switch from wave 1.5 cysts to wave 2 cysts provides a new example of developmentally programmed stem cell replacement. Previously, stem cells were shown to be capable of displacing an existing stem cell when their adhesion to the niche was greater than the adhesivity of the other resident stem cells (Jin et al. 2008). In the case of dual FSC-P replacement, neither the source of the new stem cells, how adhesivity sufficient for replacement is ensured, nor how wave 2 cysts signal replacement are known. The most plausible mechanism would be if the somatic intermingled cells surrounding early wave 2 cysts are FSC-competent and sufficiently adhesive to displace FSC-Ps and refill both niches.

### What evolutionary forces might select for similar-seeming follicle waves in Drosophila and mouse?

The general similarity of the three Drosophila follicle waves described here to those found in the mouse ovary is striking. This could result from either a coincidence, from parallel evolution, or from evolutionary conservation. Both species are sufficiently long-lived to require a major source of oocyte production to sustain adult fertility so the existence of a wave 2 is expected. Why mammals lost female GSCs, unlike lower vertebrates and many invertebrates does remain unclear. Wave 1.5 follicles provide the earliest eggs for fertility which in both species might benefit from adaptation to a subpar environment (see above). In both species, somatic cells on wave 1.5 and wave 2 follicles arise from different progenitors, which in mouse have different transcriptomes (Niu and Spradling, 2020). FSC-A transcriptomes should be compared to those of intermingled cells and adult FSCs.

The production of an initial follicle wave in both species that does not produce oocytes, but undergoes programmed turnover, is especially intriguing. It is potentially significant that in both species, wave 1 follicles develop at times when steroid hormone pulses are influencing adult development, including the programming of reproductive behavior (Trova et al. 2007; Yaniv and Schuldinger, 2016). In a wide range of species, including insects and mammals, individuals show a variety of reproductive behaviors (Emlen, 2014). Although entirely speculative at present, we favor the hypothesis that wave 1 follicles provide a way to mold reproductive behavior to fit the reproductive potential of individual animals, as signaled at a late preadult stage by both germ-cell dependent and broader physiological signals. Recent advances in mapping complete brain connectomes in Drosophila, including the relatively few circuits showing sexual dimorphism, encourages the view that these ideas can be tested experimentally (Dorkenwald et al. 2024; Schlegel et al. 2024; Berg et al. 2025).

## MATERIALS AND METHODS

### Fly Husbandry and Developmental Staging

Flies were cultured in 25°C incubators on standard cornmeal/molasses/agar media, therefore all APF hours mentioned are measured at 25°C. To collect pupa with accurate APF timing, immobile wandering third instar larva are collected, and 4 hours later only the larva that underwent pupariation are retained while those that are still larva are discarded. This method ascertains that all the remaining pupa to be examined have pupariated at the same time within a ±2 hour window. We used FlyBase (FB2025_05) to find information on nomenclature, genes, etc. (Öztürk-Çolak et al. 2024).

### Lineage Tracing Design

Lineage clones from mitotic recombination were generated using heat shock by the method of Harrison and Perrimon (1993). Females of genotype X-15-33/X-15-29; MKRS, hs-FLP/+ were generated by standard crosses. To induce flippase expression, larva/pupa at various developmental times as specified in the results were heat shocked at 37°C by immersion in a circulating water bath for 1 hour. The recombination enabled by flippase constructs a functional copy of α84Btubulin-LacZ, leading to LacZ reporter being constitutively expressed in the cell where recombination happened. The LacZ-marked clone produced by proliferation of the labeled individual cells in the ovary were subsequently imaged either 24 hours (in pupae) or 5 days (in newly eclosed adult) after heat shock. Quantification of cells in lineage clones was done in Imaris by examining across the entire Z-stack of tissue.

### Immunostaining and Light Microscopy

Drosophila pupal and adult ovaries were dissected in phosphate-buffered saline (PBS) followed by immediate fixation in 4% paraformaldehyde in PBS (made from stock 16% Paraformaldehyde Aqueous Solution EM Grade, Electron Microscopy Sciences, SKU#15710) for 30 minutes at room temperature on rocking platform. Samples were then washed and permeabilized twice with 1% PBT (Triton X-100 in PBS) at 10 minutes each, blocked in 1% PBTB (0.6% BSA and 5% NGS in 1%PBT) overnight at 4°C, and incubated with primary antibodies in 0.3% PBTB (0.6% BSA and 5% NGS in 0.3%PBT) overnight at 4°C. Next, samples were washed 2 times with 0.3% PBT and 1 time with 0.3%PBTB at 30 minutes each. Secondary antibodies were then added in 0.3% PBTB and incubation was done overnight at 4°C, then samples were washed 3 times with 0.3% PBT, the second time including 1:10,000 DAPI if nuclear stain was desired. The neutral lipid dye, BODIPY 493/503 (D3922 ThermoFisher) at 10mM, was used at 1:250 for 40’ in 0.3% PBT. Lastly, the sample was briefly washed in PBS before mounting on slide in Vectashield Antifade mounting medium. All overnight incubations were performed on a platform rocker and other shorter incubations on a nutator.

Primary antibodies used were GFP (1:200, chicken, Abcam Ab13970), LacZ (1:200, chicken, Abcam Ab134435), vasa (1:50,000, rabbit, from Dr. Ruth Lehmann) and 1B1/hts (1:50, mouse, from Developmental Studies Hybridoma Bank). Secondary antibodies used were Invitrogen AF-488, AF-568 and AF-680, all at 1:250. Images were acquired on Leica TCS SP8 confocal microscope, using 10X/NA0.25 objective for ovarioles and 63X/NA1.4 objective for germaria and pupal ovaries.

### Image Analysis and Cell Quantification

Z-stack images acquired were re-constructed into 3D object using Imaris software (v10.0.0). Statistical analysis of cells was conducted using the spot function with manual spot assignment and automated quantification. Lineage tracing clone size was determined based on the number of nls-lacZ+ nuclei, and PGC/cyst number in the pupal ovaries was determined based on the number of hts+ spectrosomes/fusomes (sample shown in Supplemental Movie S2, a 6h APF pupal ovary with white circles showing spots assigned to each spectrosome/fusome). All data from the five induction times are reported in Table S3.

### Lineage clone quantifications

Figures 2C, 2G, 2H, 5A, 5E, and 5I are quantifications of lineage clones taken from the full dataset recorded in Supplemental Table S3. Figures 2C, 5A or by the count of a given clone type out of all ovarioles scored (Figures 5E, 5I). Labeling rates were calculated by dividing the number of ovarioles containing the given clone type over the total number of ovarioles. Clone counts were the raw number of total clones observed of a given type, standard errors are inapplicable since these are single numbers. Figure 2H shows the percentage of wave 1.5 cysts at each time point with 8, 4, 2 or 1 labeled germ cells, corresponding to recombination in a CB/differentiating PGC, 2CC, 4CC or 8CC respectively (see Figure 1E). These characteristics are fixed at the time of recombination for cells committed to (PGC/CB) or undergoing cyst formation, regardless of the subsequent kinetics of cyst completion; all wave 1.5 cysts have completed cyst divisions by eclosion. Figure 2H is a plot with least square regression fit of the labeling rate change vs time of the clone categories. Figure 2G shows that PGCs that have not yet entered cysts division, do so preferentially at the 10h time point in the earliest wave 1.5 follicles, using the same data as in Figure 2H.

## Data availability

Complete results from the lineage studies are included in the paper and Supplemental material. Original Z-stack images are available upon request.

## Acknowledgements

ACS is an Investigator of the Howard Hughes Medical Institute which provided funding for this study. The authors are grateful to the Carnegie Institution for Science for hosting the Spradling laboratory and for long term support of its research. We thank Bhawana Maurya and Qi Yin for discussions and many other members of the Spradling lab for useful comments on the manuscript. We also thank Mahmud Siddiqi for microscopy and software technical support.

