## Supplementary material for "The Drosophila ovary produces two waves of adult follicles and a novel pupal wave that turns over": 687893Supplement

**The supporting information includes:**

Supplemental Table S1 to S2
Supplemental Results
Supplemental Figures S1 to S4
Legends for Supplemental Figures S1 to S4
Legends for Movies S1 to S4

Other supporting materials for this manuscript include the following:

Supplemental Table S3.xlsx

Movies S1 toS4

Supplemental Tables

Table S1

| | | GSC | R2a | R2b | | Stage 1 | | Stage 2-3 | | Stage 4-5 | | Stage 6-7 | | Stage 8-9 | | Stage 10A | |
|---|---|---|---|---|---|---|---|---|---|---|---|---|---|---|---|---|---|
| | **Ovarioles** | **16** | **16** | **4-8** | **16** | **4-8** | **16** | **2-8** | **16** | **2-8** | **16** | **1-8** | **16** | **1-8** | **16** | **1-8** | **16** |
| **-2h** | **346** | 70 | 13 | 6 | 33 | 24 | 9 | 37 | 6 | 34 | 5 | 59 | 4 | 60 | 1 | 2 | 0 |
| **2h** | **303** | 59 | 34 | 3 | 51 | 25 | 8 | 51 | 11 | 65 | 7 | 65 | 3 | 81 | 0 | 67 | 0 |
| **6h** | **129** | 37 | 3 | 2 | 15 | 11 | 2 | 15 | 2 | 28 | 1 | 21 | 1 | 48 | 0 | 49 | 0 |
| **10h** | **224** | 65 | 14 | 18 | 33 | 37 | 4 | 51 | 5 | 56 | 4 | 88 | 0 | 116 | 1 | 24 | 0 |
| **14h** | **164** | 45 | 13 | 17 | 22 | 27 | 4 | 39 | 5 | 56 | 3 | 85 | 1 | 107 | 0 | 1 | 0 |
| **Total** | **1166** | **276** | **77** | **46** | **154** | **124** | **27** | **193** | **29** | **239** | **20** | **318** | **9** | **412** | **2** | **143** | **0** |

| | GSC | R2a | R2b | | Stage 1 | | Stage 2-3 | | Stage 4-5 | | Stage 6-7 | | Stage 8-9 | | Stage 10A | |
|---|---|---|---|---|---|---|---|---|---|---|---|---|---|---|---|---|
| | **16** | **16** | **4-8** | **16** | **4-8** | **16** | **2-8** | **16** | **2-8** | **16** | **1-8** | **16** | **1-8** | **16** | **1-8** | **16** |
| **-2h** | 18.2% | 3.8% | 1.7% | 9.5% | 6.9% | 2.6% | 10.7% | 1.7% | 9.8% | 1.4% | 17.1% | 1.2% | 17.3% | 0.3% | 0.6% | 0% |
| **2h** | 18.0% | 11.2% | 1.0% | 16.8% | 8.3% | 2.6% | 16.8% | 3.6% | 21.5% | 2.3% | 21.5% | 1.0% | 26.7% | 0.0% | 22.1% | 0% |
| **6h** | 17.8% | 2.3% | 1.6% | 11.6% | 8.5% | 1.6% | 11.6% | 1.6% | 21.7% | 0.8% | 16.3% | 0.8% | 37.2% | 0% | 38.0% | 0% |
| **10h** | 28.9% | 6.3% | 8.0% | 14.7% | 16.5% | 1.8% | 22.8% | 2.2% | 25.0% | 1.8% | 39.3% | 0% | 51.8% | 0.4% | 10.7% | 0% |
| **14h** | 28.3% | 7.9% | 10.4% | 13.4% | 16.5% | 2.4% | 23.8% | 3.0% | 34.1% | 1.8% | 51.8% | 0.6% | 65.2% | 0% | 0.6% | 0% |

Table S1. Germ cell clone quantification. The table on top records the number of germ cell clones at different sizes, stages and HS time points, with the total number of ovarioles in the first column. The bottom table shows lineage labeling frequency (number of clones/number of total ovarioles) of all clone types in the top table. The first header row indicates the location of recorded germline clones, and the second header row indicates the size of such clones (e.g. 4-8 represents tally of 4-cell and 8-cell germline clones, etc.) For germ cells, only one replicate group is fully quantified due to low level of variation (n=346, 303, 129, 224, 164 ovarioles for each time point, respectively).

Table S2

| | Ovarioles | 1CC | 2CC | 1EC | 2EC | EC-M | FSC-A | FSC-P (with FSC) | Long FSC-P (without FSC) | Short FSC-P (without FSC) |
|---|---|---|---|---|---|---|---|---|---|---|
| -2h-1 | **346** | | | 2 | 2 | 1 | 6 | 5 | 5 | 13 |
| -2h-2 | **275** | 1 | | 2 | 3 | 1 | 1 | 2 | 2 | 6 |
| 2h-1 | **303** | 4 | 1 | 5 | | | 7 | 2 | | 6 |
| 2h-2 | **214** | 3 | | 4 | 5 | | 7 | 1 | 1 | 7 |
| 6h-1 | **129** | 1 | | 4 | | | 5 | 1 | | 13 |
| 6h-2 | **320** | 5 | 1 | 12 | 4 | 2 | 7 | 2 | 3 | 16 |
| 10h-1 | **224** | 19 | 7 | 24 | 11 | 17 | 17 | 10 | 6 | 52 |
| 14h-1 | **164** | 10 | 1 | 10 | 2 | | 11 | 1 | 11 | 32 |
| 14h-2 | **111** | 4 | | 7 | 5 | 1 | 6 | 6 | 5 | 9 |
| 14h-3 | **184** | 9 | 1 | 23 | 14 | 1 | 13 | 7 | 7 | 29 |

| | 1CC | 2CC | 1EC | 2EC | EC-M | FSC-A | FSC-P (with FSC) | Long FSC-P (without FSC) | Short FSC-P (without FSC) |
|---|---|---|---|---|---|---|---|---|---|
| -2h | 0.36% | 0% | 0.65% | 0.83% | 0.33% | 1.05% | 1.09% | 1.09% | 2.97% |
| 2h | 1.36% | 0.33% | 1.76% | 2.34% | 0% | 2.79% | 0.56% | 0.47% | 2.63% |
| 6h | 1.17% | 0.31% | 3.43% | 1.25% | 0.63% | 3.03% | 0.70% | 0.94% | 7.54% |
| 10h | 8.48% | 3.13% | 10.71% | 4.91% | 7.59% | 7.59% | 4.46% | 2.68% | 23.21% |
| 14h | 4.86% | 0.58% | 8.30% | 4.44% | 0.72% | 6.39% | 3.27% | 5.01% | 14.46% |

Table S2. Somatic cell clone quantification. The top table is the raw count of each type of somatic cell clone at each HS time point, the bottom table shows the labeling rates in percentage of each type of somatic cell clone (number of ovarioles with such clone over the total number of ovarioles). The top table shows the count of each replicate (e.g. 2h-1 and 2h-2 are two replicates of the 2h HS time point), while the bottom table shows the average labeling rate of each time point. 1CC: clone of 1 cap cell; 2CC: clone of 2 cap cells; 1EC: clone of 1 EC; 2EC: clone of 2 ECs; EC-M: clone with 4 or more ECs (87% are 4 EC clones); FSC-A: number of short adult FSC clones that correspond to wave 2 follicles (see text); FSC-P (with FSC): number of long FSC clones that contains an active pupal FSC (see text); long FSC-P (without FSC): number of pupal FSC clones covering multiple consecutive wave 1.5 follicles that are missing an active FSC (see text); short FSC-P (without FSC): number of short pupal FSC clones covering only 1 or 2 of the most mature wave 1.5 follicles (see text). n=621, 517, 449, 224, 459 total scored ovarioles for each time point, respectively.

#### External Supplemental Tables

Table S3.xlsx. Raw data recording the location and size of each individual observed clone. For interpretation please view the first legends tab inside the excel file.

#### Supplemental Results

<u>The first wave 1.5 follicles have developmental abnormalities</u>

Follicular coverage of the first wave 1.5 follicles in each ovariole included contributions from a different source in addition to FSC-Ps. Terminal wave 1.5 follicle cell clones frequently contain labeled follicle cells in a posterior patch and less often in other positions (Figure S3G-G'). Basal stalk cells were previously reported to contribute to pupal follicle cells (Reilein et al., 2021), and movement of basal stalk cells onto the posterior wave 1.5 follicle layer provides an attractive explanation for the bias terminal follicle clones show toward the posterior region where the stalk attaches. Similar behavior is also documented in the adult ovarioles, where inter-follicular stalk cells are known to sometimes offload extra cells onto the adjoining follicle (Nystul and Spradling, 2010).

The oldest 1-2 wave 1.5 follicles also manifested small developmental changes compared to normal adult follicles. They contain oversized polar cells, suggesting increased polyploidization (Figure S2F). The number and size of border cell clusters, which include the anterior polar cells were sometimes enlarged and delayed in migration (Figure S2F-G). These wave 1.5 follicles also have a high rate of germline cyst developmental abnormality where they contain 8 or 32 cells per follicle (Figure S2H), indicating they underwent one less or one more round of cyst division.

Supplemental Figures S1 to S4

**Figure S1.** Additional Information on Experimental Design and PGC biology

**A.** Schematic showing the detailed recombination process that gives rise to a lineage-labeled cell. **B.** An asymmetry in ovariole formation and PGC distribution exists in early pupal ovary, most prominent at 2h-6h APF and progressively becomes less evident over time. No significant asymmetry is observed by 18h APF. Ovarioles on the top of the image include more germ cells than the ovarioles on the bottom, as indicated by the brackets. PGCs posterior to the ovarioles are also unevenly distributed, more enriched on one side. 3D view shown in Movie S1. **C.** Example ovariole at 30h APF and 40h APF, both lacking the adult 2a/2b junction and R2b cyst morphology. **D-E.** Summary diagrams showing the labeling status of germ cells in wave 1.5 follicles following induction at -2h (F), or 14h (G).

**Figure S2.** Additional Quantifications of Wave 1 germ cells

**A.** Quantification of PGCs and cysts in the entire pupal ovary from -2h to 30h APF, showing the extent of PGC symmetric division and the dip caused by wave 1 loss starting at 22h APF. Quantification is based on counting the number of spectrosomes/fusomes, example counting process shown in Movie S2. **B.** Images showing the last rounds of PGC division take place among PGCs that give rise to wave 1.5 follicles at 14h APF. These are the PGCs that become included in the forming ovarioles but are too posterior to become GSCs. No more such divisions are observed at any time point later than 14h APF. **C.** A zoomed-out image of Figure 3N', showing that the single germ cells with vacuole inclusion are indeed GSCs next to the terminal filament, appearing as stacks of DAPI+ flat cells to the left side of the image. **D.** Quantification of wave 1 germ cells (red) and cysts (blue) from 14h to 22h APF. Germ cell count stays the same while cyst number drops from 18h to 22h APF, due to the loss of wave 1 cysts initiating at 22h while the overall number being compensated by continuous cyst division. **E.** Zoomed-in display of the blue "Total Cysts" bars in S2A, detailing the relative proportion of cysts at different sizes in the total population. **F.** A final wave 1.5 follicle now at stage 9 that was lineage labeled at 14h and contains a follicle cell clone which extends to the previous follicle. The border cell cluster at the anterior is mosaic (magnified in the box below) with 3 marked cells, and migrates slower than in adults toward the posterior. Other examples show the large size of the polar cells in these clusters and the presence of 3-4 labeled cells. **G-G'.** More examples showing polar cells migrate posteriorly at different speed than main body follicle cells, while in adults they migrate together. **H-H'''.** Ovarioles of newly eclosed flies showing abnormal cyst size of 8 cells and 32 cells in the last two wave 1.5 follicles, and the disproportionate polyploidization of several nurse cells.

**Figure S3.** Additional examples of FSC clones

**A-A'.** Ovarioles with long FSC-P clones that retained an active pupal FSC. **B-C.** Examples of FSC-A clones. C' is the magnified germarium region of the ovariole in C. **D-D'.** Ovarioles with long FSC-P clones that do not contain an active pupal FSC, replaced by an unlabeled FSC. **E-E'.** Short FSC-P clones on the older wave 1.5 follicles that had an early FSC loss. **F-F'.** The smallest follicle cell clones that only cover the anterior portion of the most mature wave 1.5 follicle, complementing the posterior patch pattern shown in G-G'. **G-G'.** Small patches of follicle cell clone that only cover the most mature wave 1.5 follicles, most likely originating from the basal stalk cells as previously reported (Reilein et al. 2021).

**Figure S4.** Lineage labeled escort cell (EC) clones.

**A.** A germarium stained to mark the ECs using the enhancer trap line PZ1444. Two GSCs (unlabeled) are seen at the front associated with cap cells (arrowhead). In addition, 8 specific ECs are labeled and indicated with arrowheads and assigned corresponding names. B-L show germaria following clonal marking to identify EC-specific clones. **B-B''.** Example germaria each with one labeled EC1 (arrows). **C-C'.** Germaria with single EC2 clone. **D.** Germarium with an EC3 clone. **E-E'.** Germaria with single EC5 clone. **F-F'.** Examples of EC6 clone, immediately next to FSCs. **G.** An EC6 clone in the middle of the germarium at 2a/2b junction. **H.** Labeling frequency of EC clones based on the number of ECs each clone contains. **I.** Frequency of ovarioles with clones of ECs by type (EC1-6). **J.** Two adjacent labeled ECs, E5 and E4 are located directly adjacent to an unlabeled FSC. **J'.** An ovariole containing 4 ECs that are close to the 2a/2b junction. **K-K''.** Examples of cap cell clones. K' contains 2 labeled cap cells. **L.** An ovariole containing both a CB clone and a FSC-P clone that has lost its active pupal FSC, showing FSC-P loss happened slightly after the CB clone passed by 2a/2b junction.

Legends for Movies S1 to S2

Movie S1: Rotation of a 6h APF ovary showing the asymmetry of PGC distribution along a dorsal-ventral axis.

Movie S2: Example process of PGC counting in the pupal ovary. Spectrosomes/fusomes shown as magenta dots stained by Hts/1B1 are circled, as they are counted in Imaris using the Spot function.

Movie S3-S4: Examples showing how the quantification of follicle cell coverage in Figure 5K-L was done in Imaris in 3D reconstruction. Movie S3 shows an FSC-L clone and Movie S4 shows an FSC-M clone.

### Figure S1

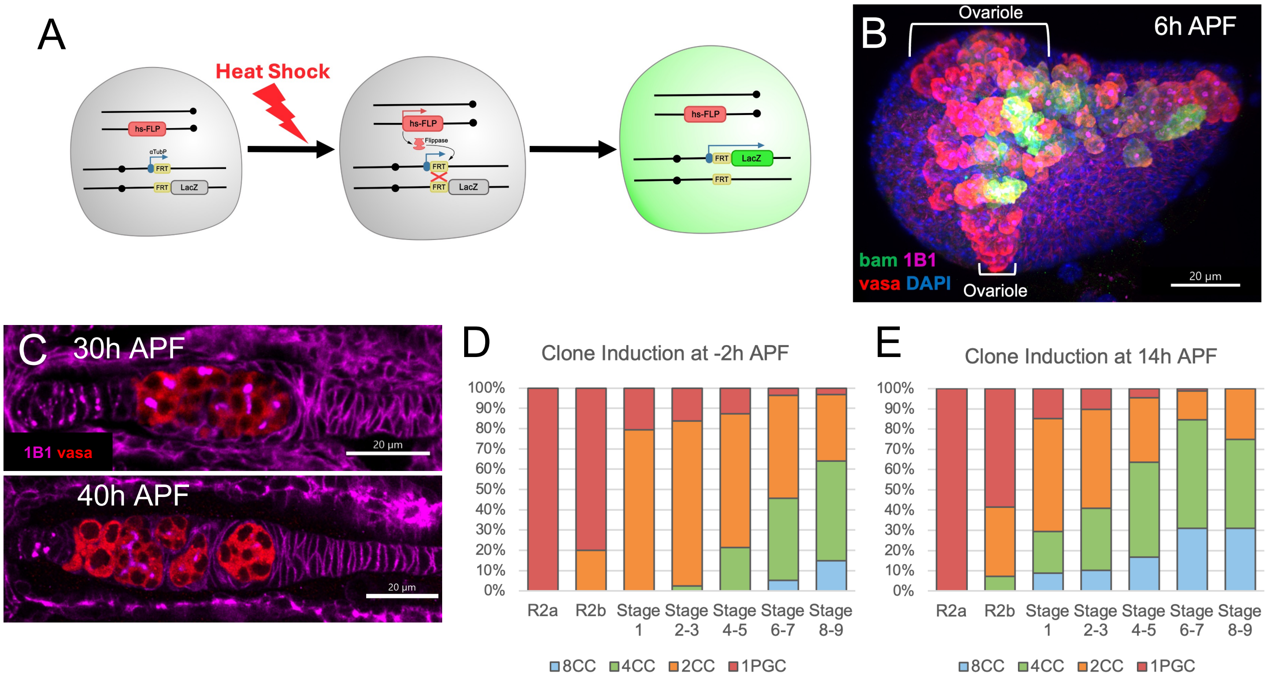
A
Heat Shock
hs-FLP
αTubP
FRT
LacZ
Flippase
B
Ovariole
6h APF
bam 1B1
vasa DAPI
Ovariole
20 μm
C
30h APF
1B1 vasa
20 μm
40h APF
20 μm
D
Clone Induction at -2h APF
100%
90%
80%
70%
60%
50%
40%
30%
20%
10%
0%
R2a
R2b
Stage 1
Stage 2-3
Stage 4-5
Stage 6-7
Stage 8-9
8CC
4CC
2CC
1PGC
E
Clone Induction at 14h APF
100%
90%
80%
70%
60%
50%
40%
30%
20%
10%
0%
R2a
R2b
Stage 1
Stage 2-3
Stage 4-5
Stage 6-7
Stage 8-9
8CC
4CC
2CC
1PGC

### Figure S2

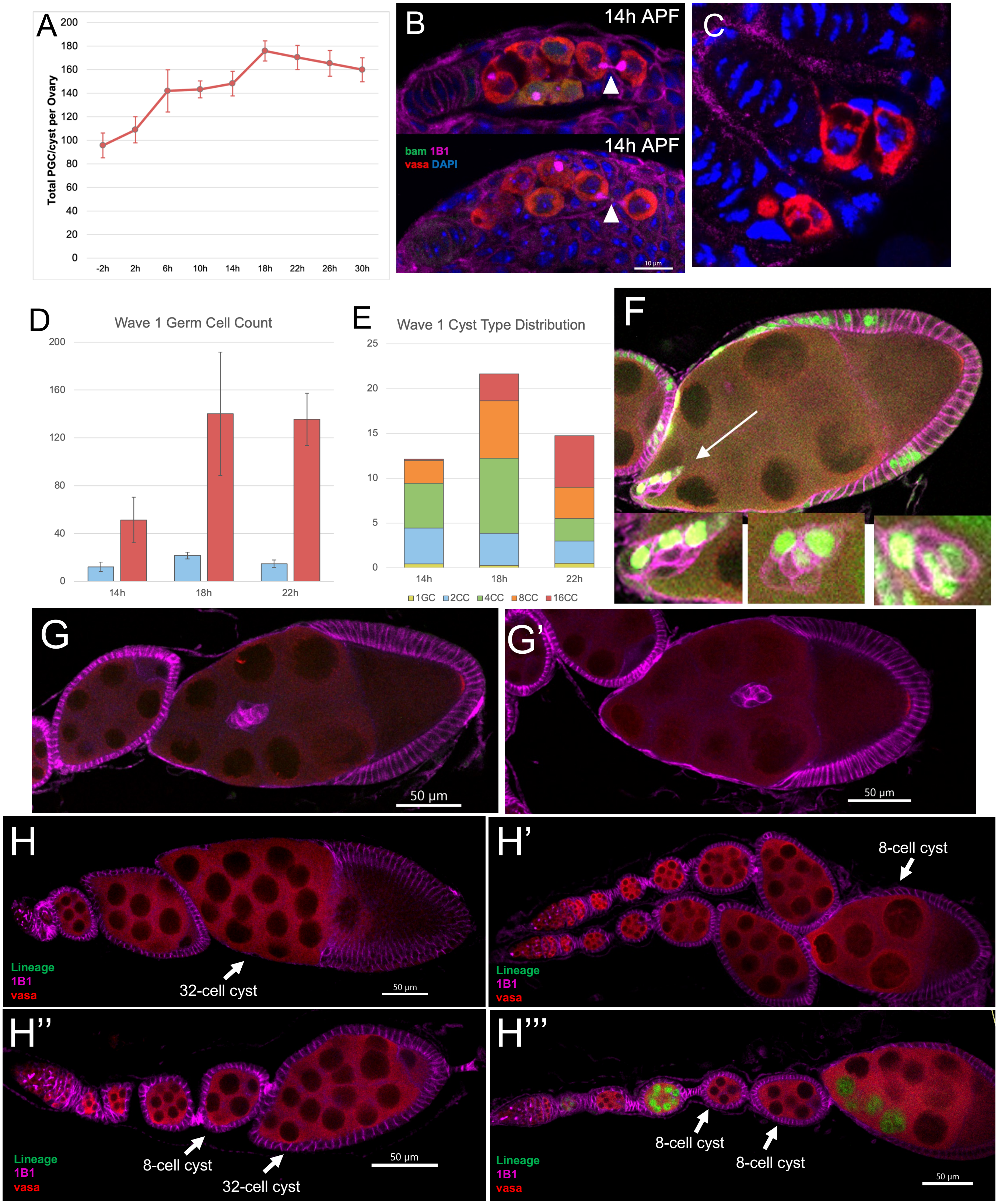

A
Total PGC/cyst per Ovary
200
180
160
140
120
100
80
60
40
20
0
-2h
2h
6h
10h
14h
18h
22h
26h
30h
B
14h APF
14h APF
bam 1B1
vasa DAPI
10 µm
C
D
Wave 1 Germ Cell Count
14h
18h
22h
E
Wave 1 Cyst Type Distribution
1GC 2CC 4CC 8CC 16CC
F
G
50 µm
G'
50 µm
H
Lineage
1B1
vasa
32-cell cyst
50 µm
H'
8-cell cyst
Lineage
1B1
vasa
H''
Lineage
1B1
vasa
8-cell cyst
32-cell cyst
50 µm
H'''
8-cell cyst
8-cell cyst
Lineage
1B1
vasa
50 µm

### Figure S3

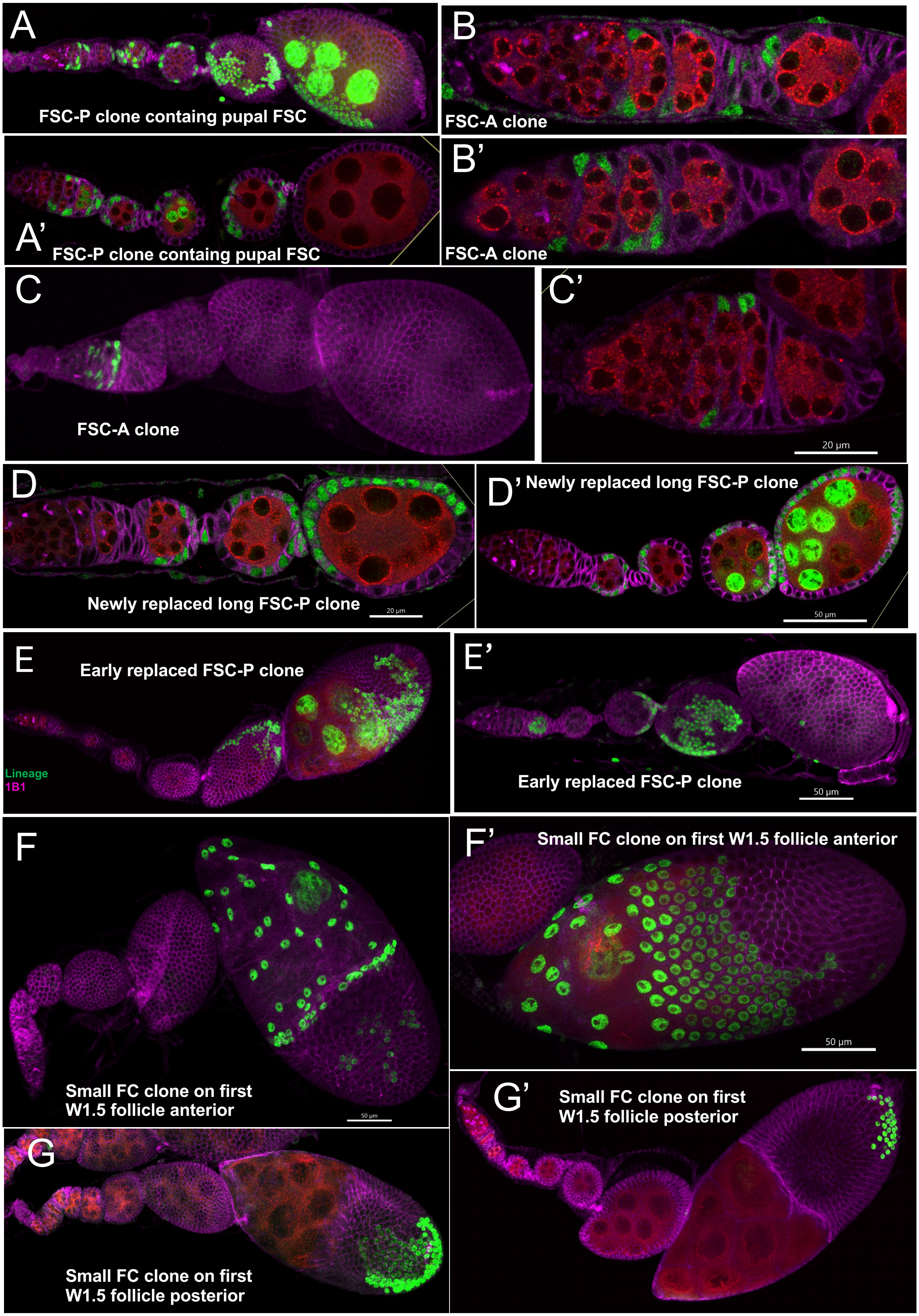
A
FSC-P clone containg pupal FSC
A'
FSC-P clone containg pupal FSC
B
FSC-A clone
B'
FSC-A clone
C
FSC-A clone
C'
20 µm
D
Newly replaced long FSC-P clone
20 µm
D' Newly replaced long FSC-P clone
50 µm
E
Early replaced FSC-P clone
Lineage
1B1
E'
Early replaced FSC-P clone
50 µm
F
Small FC clone on first
W1.5 follicle anterior
50 µm
F' Small FC clone on first W1.5 follicle anterior
50 µm
G
Small FC clone on first
W1.5 follicle posterior
G'
Small FC clone on first
W1.5 follicle posterior

### Figure S4

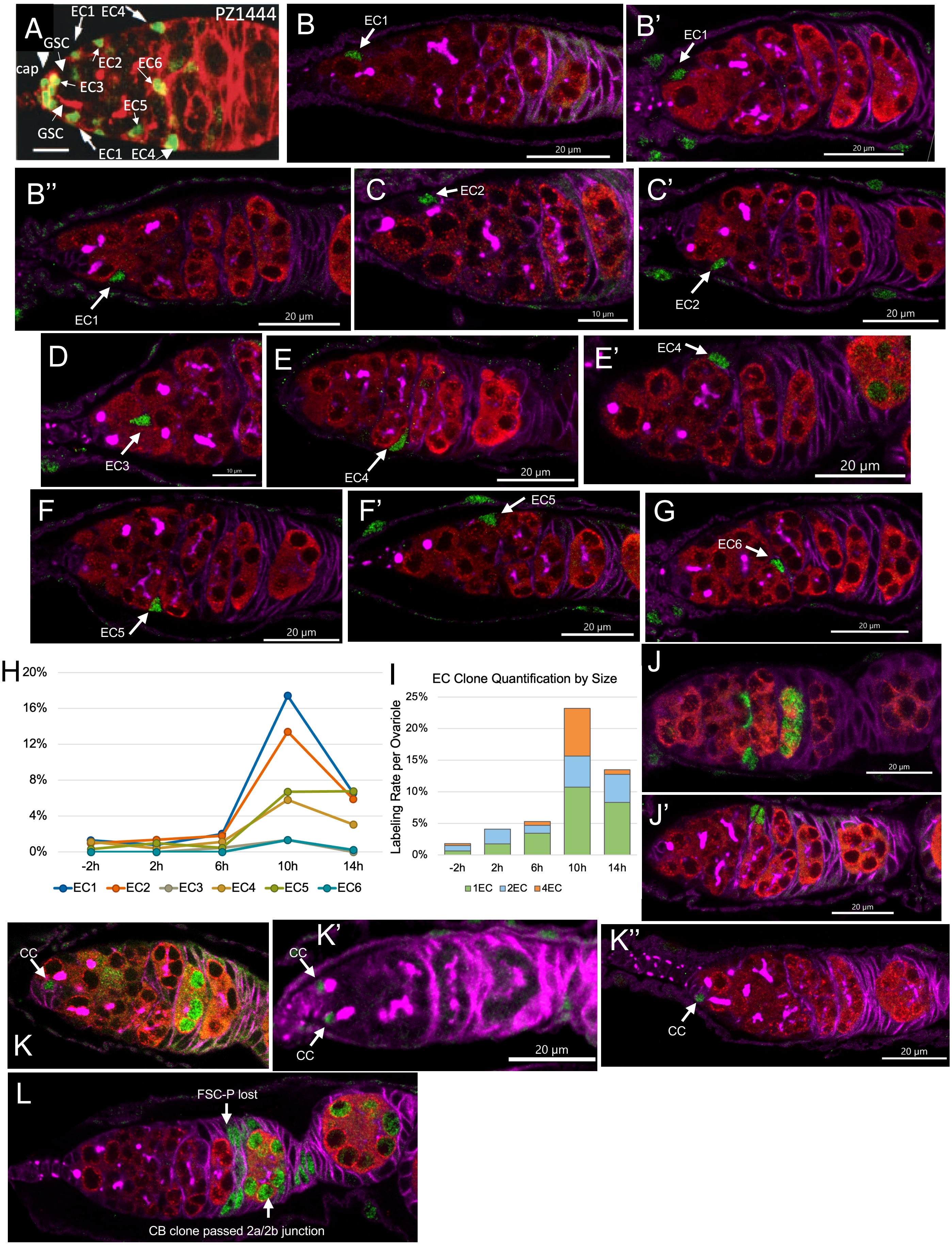

A
PZ1444
EC1 EC4
GSC
cap
EC2
EC6
EC3
EC5
GSC
EC1 EC4
B
EC1
20 µm
B'
EC1
20 µm
B''
EC1
20 µm
C
EC2
10 µm
C'
EC2
20 µm
D
EC3
10 µm
E
EC4
20 µm
E'
EC4
20 µm
F
EC5
20 µm
F'
EC5
20 µm
G
EC6
20 µm
H
20%
16%
12%
8%
4%
0%
-2h
2h
6h
10h
14h
EC1
EC2
EC3
EC4
EC5
EC6
I
EC Clone Quantification by Size
Labeling Rate per Ovariole
25%
20%
15%
10%
5%
0%
-2h
2h
6h
10h
14h
1EC
2EC
4EC
J
20 µm
J'
20 µm
K
CC
K'
CC
CC
20 µm
K''
CC
20 µm
L
FSC-P lost
CB clone passed 2a/2b junction
